# Environmental DNA metabarcoding for biodiversity monitoring of a highly-diverse tropical fish community in a coral-reef lagoon: Estimation of species richness and detection of habitat segregation

**DOI:** 10.1101/2020.05.07.083261

**Authors:** Shin-ichiro Oka, Hideyuki Doi, Kei Miyamoto, Nozomi Hanahara, Tetsuya Sado, Masaki Miya

## Abstract

An environmental DNA (eDNA) metabarcoding approach has been widely used for biodiversity monitoring of fishes, although it has rarely been applied to tropical and subtropical aquatic ecosystems, where species diversity is remarkably high. This study examined the extent to which species richness can be estimated in a small coral reef lagoon (1500 × 900 m) near Okinawa Island, southern Japan, where the surrounding waters are likely to harbor more than 1500 species of fish. During 2015–2017, a total of 16 capture-based surveys were conducted to create a faunal list of fish species, followed by eDNA metabarcoding based on seawater samples taken from 11 sites in the lagoon on a day in May 2019. We also tested whether eDNA metabarcoding could detect differences between adjacent fish communities inhabiting the offshore reef edge and shore-side seagrass beds within the lagoon. A total of 217 fish species were confirmed by the capture-based samplings, while 291 fish species were detected by eDNA metabarcoding, identifying a total of 410 species distributed across 119 families and 193 genera. Of these 410 species, only 96 (24% of the total) were commonly identified by both methods, indicating that capture-based surveys failed to collect a number of species detected by eDNA metabarcoding. Interestingly, two different approaches to estimate species richness based on eDNA data yielded values close to the 410 species, including one that suggested an additional three or more eDNA surveys from 11 sites (36 samples) would detect 90% of the 410 species. In addition, non-metric multi-dimensional scaling for fish assemblages clearly distinguished between the fish communities of the offshore reef edge and those of the shore-side seagrass beds.

## 1 INTRODUCTION

Biodiversity monitoring is essential for ecosystem conservation and the sustainable use of biological resources (Boulinier, Nichols, Sauer, Hines, & Pollock et al., 2019; Lepetz, Massot, Schmeller, & Clobert, 2009; Reimer et al., 2019). In case of fishes, traditional biodiversity monitoring has been conducted through direct capture of specimens and/or underwater visual observations (Evans et al., 2017; Pelletier, Leleu, Mou-Tham, Guillemot, & Chabanet, 2011; Thomsen et al., 2012). The former approach is invasive and requires subsequent morphological identification of specimens based on taxonomic expertise, while the latter is not invasive, although visual identification by observers with different skill levels involves great uncertainty, as they are likely to overlook rare or elusive species (Brock, 1982; Edgar, Barrett, & Morton, 2004; Harvey, Fletcher, Shortis, & Kendrick, 2004). In addition, both approaches are costly and time-consuming, and their feasibility heavily depends on weather and water conditions (Andruszkiewicz et al., 2017; Evans et al., 2017; Sigsgaard et al., 2017; Thomsen & Willerslev, 2015). Therefore, it is practically impossible to perform continuous biodiversity monitoring of fish communities at multiple sites using traditional approaches.

Environmental DNA (eDNA), which comprises indirect genetic markers of macroorganisms such as feces, body mucus, blood, and sloughed tissue or scales, has emerged as an alternative (Kelly, Port, Yamahara, & Crowder, 2014). In particular, the eDNA metabarcoding (EDM) approach enables simultaneous detection of multiple species using a high-throughput next-generation sequencing (NGS) platform (e.g., Taberlet, Coissac, Hajibabaei, & Rieseberg, 2012; Kelly, Port, Yamahara, Martone, et al., 2014, Miya et al. 2015). This approach amplifies a short fragment of eDNA from the target taxa (e.g., fishes) using a set of universal PCR primers and appends various adapter and index sequences to both ends of the amplified fragments (amplicons). Various combinations of different index sequences enables massive parallel sequencing using the NGS platform, with an output comprising tens of millions of amplicon sequences from multiple sampling sites. After data preprocessing and subsequent taxon assignment using a bioinformatics analysis pipeline, a tentative list of species is available for each sampling site.

Owing to its cost-effectiveness and high sensitivity for the detection of fish species, EDM has been applied to various aquatic environments such as rivers (McDevitt et al., 2019; Morita et al., 2019; Nakagawa et al., 2018; Sales, Wangensteen, Carvalho, & Mariani, 2019), lakes (Fujii et al., 2019; Lawson Handley et al., 2019), brackish (Zhang, Yoshizawa, Iwasaki, & Xian, 2019; Zou et al., 2020), coastal (Andriyono, Alam, & Kimet al., 2019; Andriyono et al., 2019; Andruszkiewicz et al., 2017; Yamamoto et al., 2017) and deep waters (Thomsen et al., 2016) across temperate latitudes with moderate species richness. These studies have successfully detected diverse fish types in their respective ecosystems and demonstrated that EDM often outperforms other inventory methods such as capture-based sampling (CBS) and visual observations. However, few EDM studies have been conducted on fish in tropical and subtropical coastal ecosystems (Andriyono et al., 2019; Nguyen et al., 2020; Sigsgaard et al., 2019), there has been no such study in coral reef lagoons, where species richness is remarkably high (Komyakova et al., 2013) [but see Miya et al. (2015) for a preliminary eDNA metabarcoding experiment].

The first objective of this study was to test the feasibility of EDM for biodiversity monitoring of highly diverse tropical and subtropical coastal fishes. Therefore, we estimated species richness in a coral reef lagoon on Bise, Okinawa Island, southern Japan, which was chosen as a test field because 16 capture-based surveys were performed there prior to this study and a list of collected fish species was available. In addition to these two datasets, we also conducted a literature survey to create a faunal list of fish species that are likely to be found in the lagoon. By comparing the results of EDM and CBS with the faunal list, we evaluated the efficiency and reproducibility of both methods.

The second objective was to test the sensitivity of EDM for detecting habitat segregation in fish communities within the lagoon by comparing those in the offshore reef edge and the shore-side seagrass beds. Coral reefs and seagrass beds are often located adjacent to one another (Dorenbosch, Grol, Nagelkerken, & van der Velde et al., 2005) and the coral reef lagoon in Bise was no exception. However, fish communities change along a coral reef-seagrass bed gradient (Dorenbosch et al., 2005; Fabricius, De’ath, McCook, Turak, & Williams, 2005). Therefore, we tested the sensitivity of EDM to determine if it could detect differences in fish community compositions between offshore reef edges and on shore-side seagrass beds. In addition, we tested the differences in the ecological and biological traits of fish, such as body size and shape, trophic position, and habitat preference, between the species frequently detected by eDNA from offshore and shore-side sites.

To validate the feasibility and sensitivity of EDM, we used MiFish primers (Miya et al. 2015) for library preparation before NGS analysis, which have been demonstrated to outperform other competing primers in recent studies (Bylemans et al., 2018; Collins et al., 2019) and have actually been used in various aquatic environments in and around six continents (Andruszkiewicz et al., 2017; Bylemans et al., 2018; Mariani, Baillie, Colosimo, & Riesgo, 2019; McDevitt et al., 2019; Morita et al., 2019; Nakagawa et al., 2018; Sales et al., 2019; Yamamoto et al., 2017; Zhang et al., 2019; Zou et al., 2020).

## 2 MATERIALS AND METHODS

### 2.1 Ethics statement

All 16 capture-based surveys and water sampling at the 11 sites were conducted in compliance with Japanese and local laws and regulations.

### 2.2 Capture-based sampling and literature survey

Prior to the eDNA survey, a total of 16 faunal surveys were conducted by direct capture of fish specimens from December 2015 to November 2017 in a coral reef lagoon in Bise, Motobu, Okinawa Island, Japan (Figure 1, 26°42’N, 127°52’E). The lagoon is flat and shallow, protected by the outer reef, approximately 1500 m × 900 m wide (around 1.1 km^2^). Its largest tidal amplitude during the spring tide is around 2 m, and the outer reefs are exposed at low tide, while its maximum depth reaches approximately 3 m at high tide. Coral assemblages are widely distributed in patches and eelgrass beds spread over the shore-side shallow area of the lagoon. The bottom is dominated by sand and gravel, with no distinct trace of mud.

**FIGURE 1.**
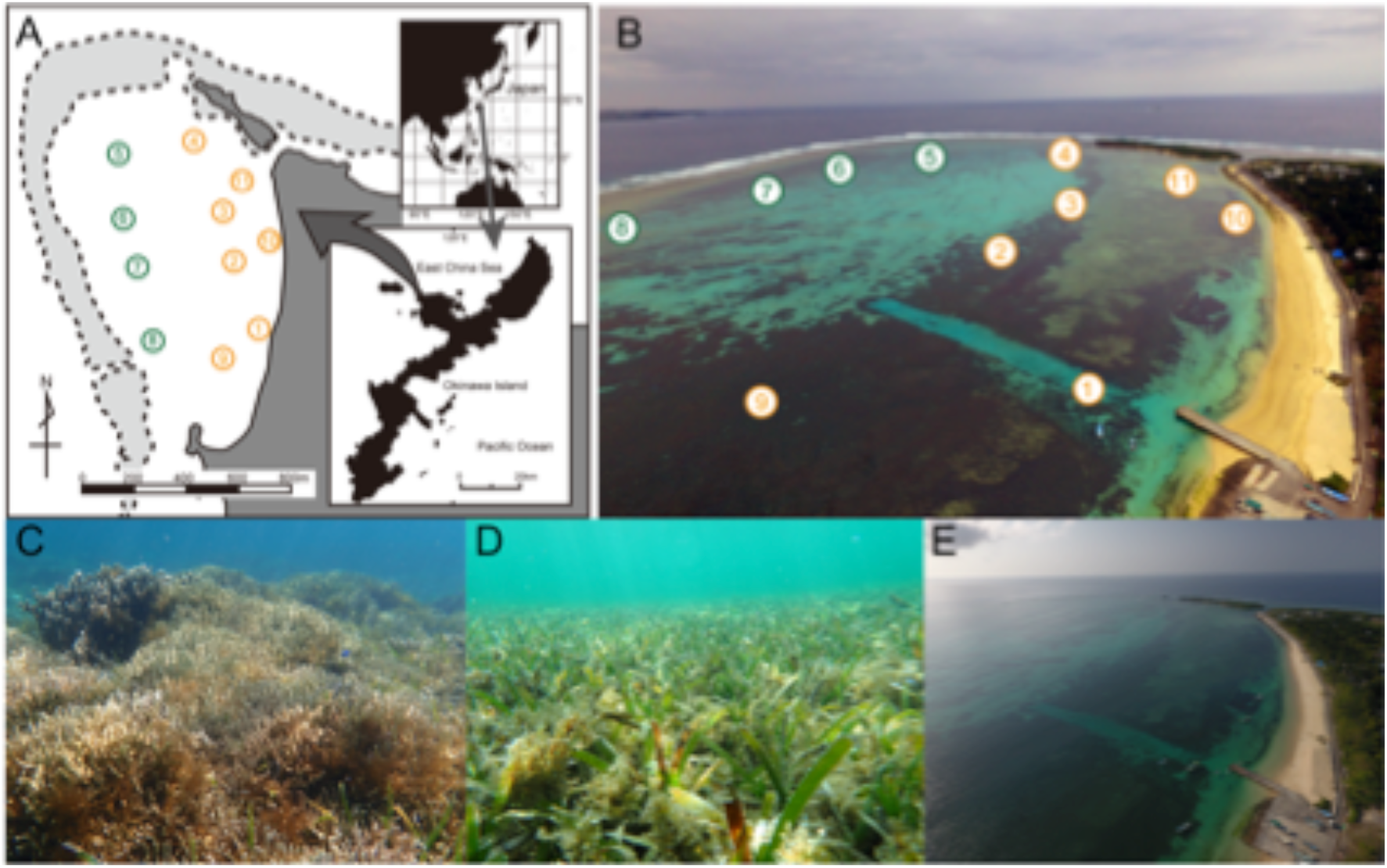
Map (A) and photo (B) showing positions of the 11 seawater sampling stations in the coral reef lagoon at Bise, Okinawa, southern Japan. The 11 samples were taken on May 8, 2019 at a tide level of 180 cm above the lowest tide near high tide. Sts. 5-8 were located along the offshore reef edge, while Sts. 1-4 and 9-11 were located on the shore-side seagrass beds. The photo (B) was taken on January 9, 2019 at a tide level of 100 cm using a remotely-operational aerial vehicle to show the position of reef edge. The lower panels show coral assemblages around St. 8 (C), eelgrass beds around St. 9 (D) and an overview of the lagoon at the same tide level of the sampling day (E). All the latter three photos were taken on May 4, 2020.

One to three researchers participated in each survey, and various combinations of capture methods, including hand, gill, and cast nettings and bait fishing, were used to collect diverse fish types (Table S1). The main purpose of the 16 surveys was to comprehensively collect fish species from the area for listing fish species and was not intended for quantitative monitoring of the fish community. For that purpose, at least one specimen in each collected fish species was deposited in the collection of the Okinawa Churashima Foundation (OCF). Those species that were difficult to capture were photographed in their habitat to provide evidence of occurrence.

In addition to the faunal survey, a literature survey was conducted to provide a baseline dataset on the faunal list of coastal fishes that are likely to occur around the Okinawa Islands. Instead of surveying primary information from the original articles, the atlas of Japanese fishes (Nakabo, 2013) was used to compile the list because it comprehensively includes the fish species recorded in Japanese waters and information has been continuously updated on the website since publication (http://www.fish-isj.jp/info/list_additon.html).

### 2.3 Water sampling and filtration

Sea water samples were collected from 11 sites in the same coral reef lagoon (Figure 1, Table 1) on 8 May 2019 using a small boat. The boat departed from a small pier near the St. 1 through the shore-side seagrass sites (Sts. 1-4), via the offshore reef-edge sites (Sts. 5-8), and finally to the most shore-side seagrass sites (Sts. 9-11). The sampling was performed during high tide to ensure the safety of boat navigation. At each site, surface seawater was collected using a disposable small cup (500 ml). The sampled water was poured into a plastic bag (DP16-TN1000, COWPACK LTD, Japan), and this process was repeated three times until the plastic bag was filled with approximately 1,000 ml seawater. Immediately after the water sampling, 1 ml benzalkonium chloride was added to the plastic bag to prevent eDNA degradation (Yamanaka et al., 2017), and the screw cap of the plastic bag was tightly closed. After water sampling, water temperature and salinity were measured with a portable water quality meter (WQC-30, DKK-TOA, Japan). Seawater samples were transported to the laboratory in a cooling box with ice packs.

**TABLE 1.** Summary of the eDNA survey at 11 stations in the lagoon along Bise, Okinawa, southern Japan on 8 May 2019

| Station | Starting time | Latitude (° N) | Longitude (° E) | Temperature (°C) | Salinity (‰) |
| --- | --- | --- | --- | --- | --- |
| Station 1 | 07:40 | 26.70356 | 127.87938 | 23.6 | 31.4 |
| Station 2 | 07:50 | 26.70581 | 127.87837 | 23.6 | 31.4 |
| Station 3 | 07:55 | 26.70756 | 127.87787 | 23.7 | 32.1 |
| Station 4 | 08:05 | 26.71001 | 127.87665 | 23.7 | 32.1 |
| Station 5 | 08:10 | 26.70932 | 127.87392 | 23.8 | 32.1 |
| Station 6 | 08:15 | 26.70700 | 127.87380 | 23.8 | 32.0 |
| Station 7 | 08:20 | 26.70573 | 127.87427 | 23.8 | 32.0 |
| Station 8 | 08:25 | 26.70298 | 127.87492 | 23.7 | 32.1 |
| Station 9 | 08:35 | 26.70230 | 127.87779 | 23.7 | 32.2 |
| Station 10 | 08:45 | 26.70656 | 127.87980 | 23.7 | 32.0 |
| Station 11 | 08:50 | 26.70862 | 127.87874 | 23.7 | 32.1 |

All water samples were filtered within 2 h of sampling in the laboratory using a combination of Sterivex filter cartridges (nominal pore size = 0.45 μm; Merck Millipore, MA, USA) and a handmade parallel filtering system using an aspirator (Figure S1). After filtration (approximately 10 min for each bag), an outlet port of the filter cartridge was sealed with parafilm, 1.6 ml RNAlater (Thermo Fisher Scientific, DE, USA) was injected into the cartridge using a disposable pipette to prevent eDNA degradation, and an inlet port was also sealed with parafilm. At the end of filtration, a filtration blank (FB) was made by filtering 1,000 ml of Milli-Q water in the same manner. The total time from the onset of water sampling to the end of filtration was about three hours.

### 2.4 DNA extraction and purification

All workspaces and equipment were thoroughly sterilized before DNA extraction. Filtered pipet tips were used, and all eDNA-extraction manipulations were conducted in a dedicated room that was physically separated from the pre- and post-PCR rooms to safeguard against cross-contamination from PCR products.

eDNA was extracted and purified following the method developed and visualized by Miya et al. (2016) and subsequently updated by Miya & Sado (2019). After vacuuming RNAlater inside the cartridge using a QIAvac 24 Plus manifold (Qiagen, Inc.), proteinase-K solution was injected into the cartridge from the unsealed inlet port, and the port was sealed again with parafilm. The eDNA captured on the filter membrane was extracted by constant stirring of the cartridge using a roller shaker placed in an incubator preheated at 56°C for 20 min. The eDNA extracts were collected from the cartridges by centrifugation and purified using a DNeasy Blood & Tissue kit (Qiagen).

### 2.5 Paired-end library preparation and MiSeq sequencing

Two-step PCR was employed to prepare paired-end libraries in the MiSeq platform (Illumina, CA, USA) and generally followed the methods developed by Miya et al. (2015) and subsequently updated by Miya and Sado (2019). For the first-round PCR (1st PCR), we used a mixture of the following four PCR primers: MiFish-U-forward (5’-ACA CTC TTT CCC TAC ACG ACG CTC TTC CGA TCT NNN NNN GTC GGT AAA ACT CGT GCC AGC-3’), MiFish-U-reverse (5’-GTG ACT GGA GTT CAG ACG TGT GCT CTT CCG ATC TNN NNN NCA TAG TGG GGT ATC TAA TCC CAG TTT G-3’), MiFish-E-forward-v2 (5’-ACA CTC TTT CCC TAC ACG ACG CTC TTC CGA TCT NNN NNN RGT TGG TAA ATC TCG TGC CAG C-3’) and MiFish-E-reverse-v2 (5’-GTG ACT GGA GTT CAG ACG TGT GCT CTT CCG ATC TNN NNN NGC ATA GTG GGG TAT CTA ATC CTA GTT TG-3’). These primer pairs co-amplify a hypervariable region of the fish mitochondrial 12S rRNA gene (around 172 bp; hereafter called MiFish sequence) and append primer-binding sites (5’ ends of the sequences before six Ns) for sequencing at both ends of the amplicon. Six random bases (Ns) in the middle of those primers were used to enhance cluster separation on the flow cells during initial base call calibrations on the MiSeq platform.

The 1st PCR was carried out with 35 cycles of a 12 μl reaction volume containing 6.0 μl 2 × KAPA HiFi HotStart ReadyMix (KAPA Biosystems, MA, USA), 1.4 μl of each MiFish primer (5 μM primer F/R), 1.2 μl sterile distilled H_2_O, and 2.0 μl eDNA template. To minimize PCR dropouts, eight technical replications were performed for the 1st PCR using a 0.2 ml 8-strips tube. The thermal cycle profile after an initial 3 min denaturation at 95°C was as follows: denaturation at 98°C for 20 s, annealing at 65°C for 15 s, and extension at 72°C for 15 s, with the final extension at the same temperature for 5 min. The 1st PCR products from the eight tubes were pooled in a single 1.5 ml tube and the pooled products were purified using a GeneRead Size Selection kit (Qiagen, Hilden, Germany) to remove dimers and monomers following the manufacturer’s protocol. Subsequently, the purified products were quantified using TapeStation 2200 (Agilent, Tokyo, Japan), and the quantified products were diluted to 0.1 ng μl-1 using Milli-Q water, which was used as a template for the second PCR.

For the second-round PCR (2nd PCR), we used the following two primers to append the dual-index sequences (8 nucleotides indicated by Xs) and flowcell-binding sites for the MiSeq platform (5’ ends of the sequences before eight Xs): 2nd-PCR-forward (5’-AAT GAT ACG GCG ACC ACC GAG ATC TAC ACX XXX XXX XAC ACT CTT TCC CTA CAC GAC GCT CTT CCG ATC T-3’); and 2nd-PCR-reverse (5’-CAA GCA GAA GAC GGC ATA CGA GAT XXX XXX XXG TGA CTG GAG TTC AGA CGT GTG CTC TTC CGA TCT-3’). The 2nd PCR was conducted with 10 cycles of a 15 μl reaction volume containing 7.5 μl 2× KAPA HiFi HotStart ReadyMix, 0.9 μl of each primer (5 μM), 3.9 μl sterile distilled H_2_O, and 1.9 μl template. The thermal cycle profile after an initial 3 min denaturation at 95°C was as follows: denaturation at 98°C for 20 s, annealing and extension combined at 72°C (shuttle PCR) for 15 s, with the final extension at the same temperature for 5 min.

To monitor contamination during the PCR process, four blank samples (negative controls) were prepared. In addition to FB, and extraction blank (EB), first and second PCR blanks (1B and 2B, respectively) with 2.0 μl Milli-Q water instead of template eDNA were prepared in the experiments.

All libraries containing the target region and the three adapter sequences were mixed in equal volumes, and the pooled libraries were size-selected from approximately 340 bp using a 2% E-Gel Size Select agarose gel (Invitrogen, CA, USA). The concentration of the size-selected libraries was measured using a Qubit dsDNA HS assay kit and a Qubit fluorometer (Life Technologies, CA, USA), diluted to 11.0 pM with HT1 buffer (Illumina, CA, USA), and sequenced on the MiSeq platform using the MiSeq v2 Reagent Kit Mini for 2× 150 bp paired-end (Illumina, CA, USA) with a PhiX Control library (v3) spike-in (expected at 5%) following the manufacturer’s protocol. All raw DNA sequence data and associated information are deposited in the DDBJ/EMBL/GenBank databases and are available from the accession number DRA009512.

### 2.6 Data preprocessing and taxon assignment

Data preprocessing and analysis of MiSeq raw reads from the 11 samples were performed using USEARCH v10.0.240 (Edgar, 2010) according to the following steps: 1) Forward (R1) and reverse (R2) reads were merged by aligning both reads using the “fastq mergepairs” command. During this process, low-quality tail reads with a cut-off threshold set at a quality (Phred) score of 2, reads too short (<100 bp) after tail trimming, and those paired reads with too many differences (>5 positions) in the aligned region (around 65 bp) were discarded; 2) primer sequences were removed from those merged reads using the “fastx truncate” command; 3) those reads without the primer sequences underwent quality filtering using the “fastq filter” command to remove low-quality reads with an expected error rate of >1% and reads too short (<120 bp); 4) the preprocessed reads were dereplicated using the “fastx uniques” command, and all singletons, doubletons, and tripletons were removed from the subsequent analyses to avoid false positives following the recommendation by the program’s author (Edgar, 2010); 5) the dereplicated reads without single-to tripletons were denoised using the “unoise3” command to generate amplicon sequence variants (ASVs) that remove all putatively chimeric and erroneous sequences (Callahan, McMurdie, & Holmes, 2017); 6) the ASVs were rarefied to the minimum read number (72,302), and 7) finally ASVs were assigned to taxon, i.e., species names (molecular operational taxonomic units; MOTUs) using the “usearch global” command with a sequence identity of >98.5% with the reference sequences (two nucleotide differences allowed) and a query coverage of ≥90%.

ASVs with sequence identities of 80%-98.5% were tentatively added U98.5 labels before the corresponding species names with the highest identities (e.g., U98.5_*Pagrus_major*) and were subjected to clustering at the level of 0.985 using the “cluster smallmem” command. An incomplete reference database needs this clustering step that enables the detection of multiple MOTUs under the same species name. Such multiple MOTUs were annotated with “gotu1, 2, 3…” and all of these outputs (MOTUs and U98.5 MOTUs) were tabulated with read abundances. Those ASVs with sequence identities of <80% (saved as “no hit”) were excluded from the above taxon assignments and downstream analyses because all of them were found to be non-fish organisms (see Results). MiFish DB ver. 36 was used for the taxon assignments, which contained 7,973 species distributed across 464 families and 2,675 genera.

To focus our analyses only on coastal fishes, all fishes principally inhabiting the coral reef lagoon were excluded. For example, deep-sea fishes (e.g., lanternfishes of the family Myctophidae), and pure freshwater fishes (e.g., members of the family Cichlidae) were excluded from further analyses.

To refine the above taxon assignments, family level phylogenies were reproduced with MiFish sequences from MOTUs, U98.5 MOTUs plus the reference sequences (contained in MiFish DB ver. 36) from those families. For each family, representative sequences (most abundant reads) from MOTUs and U98.5 MOTUs were assembled, and all reference sequences from that family were added in a fasta format. The combined fasta-formatted sequences were subjected to multiple alignment using MAFFT 7 (Katoh & Standley, 2013) with a default set of parameters. A neighbor-joining (NJ) tree was constructed with the aligned sequences in MEGA7 (Kumar, Stecher, & Tamura, 2016) using the Kimura two-parameter distances (Kimura, 1980). Distances were calculated using pairwise deletion of gaps and the among-site rate variations modeled with gamma distributions (shape parameter = 1). Bootstrap resampling (n = 100) was performed to estimate statistical support for internal branches of the NJ tree, and midpoint rooting was performed on the resulting NJ tree.

A total of 51 family level trees were visually inspected, and taxon assignments were revised as follows: For those U98.5 MOTUs placed within a monophyletic group consisting of a single genus, the unidentified MOTUs were named that genus plus sp. with sequential numbers (e.g., *Pagrus* sp. 1, sp. 2, sp. 3, …). For the remaining MOTUs ambiguously placed in the family level tree, the unidentified MOTUs were named that family plus sp. with sequential numbers (e.g., Sparidae sp. 1, sp. 2, sp. 3 …).

### 2.7 Estimation of species richness

Two approaches were employed to estimate the species richness of the study area. Statistical analyses and graphics were conducted in R ver. 3.6.0 (R Core Team, 2019) with a significance level of α = 0.05.

In the first approach, Chao II bias-corrected estimator (Chao, 2005; Olds et al., 2016) was applied to estimate the total number (Smax) of species in the survey area by EDM. Chao II estimates the minimum species richness in the system and accounts for unobserved species based on the sampling effort (n), and the number of species with one and two incidences of detection, *g* (1) and *g* (2), respectively:

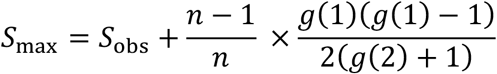

where *S*_obs_ is the total number of species directly observed. The formulation (*n* - 1) / *n* is a factor used to adjust for small sample size, which is needed for our samples of EDM (*n* = 11). The 95% confidence interval (CI) of *S*_max_ was obtained using jackknife resampling (Whitehead, 1995).

In the second approach, based on the presence/absence data of the fish species detected by eDNA, sample-based species accumulation curves were generated using “iNEXT” function in “iNEXT” ver. 2.0.19 package (Chao et al., 2014) with the first Hill number (q = 0).

### 2.8 Indicator taxa analysis

To determine which taxa had significantly different detection frequencies in the offshore reef-edge sites and shore-side seagrass beds, indicator taxa analysis (Cáceres & Legendre, 2009) was performed using the “signassoc” function of “indicspecies” package ver. 1.7.6 (Cáceres & Legedre, 2009) on the incidence-based data, with mode = 1 (group-based) and calculated the *P* values with 999 permutations after Sidak’s correction.

### 2.9 Comparisons of ecological and biological traits

The ecological and biological traits of the detected and recorded fish species were obtained from FishBase (Froese & Pauly, 2019) through searches by species names using “species” and “ecology” functions from “rfishbase” (Boettiger, Lang, & Wainwright, 2012) ver. 3.1.0 package on 10 March 2020. Differences in fish traits, body length (cm), depth range to depth (m), trophic position, preference for coral reefs, habitat preference, and lateral body shape types (categorical data) were tested for the differences between those sites along offshore reef-edge (Sts. 5-8) and on shore-side seagrass beds (Sts. 1-4 and 9-11) using general linear model (GLM) with a binomial as the error distribution. Prior to GLM, a variance inflation factor (VIF) was calculated to check the collinearity of the explanatory factors of GLMs. The maximum VIF = 1.20 was calculated, indicating that collinearity among the factors did not potentially influence the parameter estimations in GLM as VIF < 5 (Sheather, 2009).

### 2.10 Analysis of fish community structures

The differences in fish community structures were visualized using non-metric multi-dimensional scaling (NMDS) with 999 separate runs of real data. For NMDS, community dissimilarity was calculated based on incidence-based Jaccard indices, and NMDS stress was used to confirm the representation of NMDS ordination. Differences in fish community structures between sites along the offshore reef-edge (Sts. 5-8) and shore-side seagrass beds (Sts. 1-4 and 9-11) were also evaluated using permutational multivariate analysis of variance (PERMANOVA) with 999 permutations. An incidence-based Jaccard similarity matrix was used for PERMANOVA. The “metaMDS” and “adonis” functions of “vegan” ver. 2.5.6 package (Oksanen et al., 2019) were used for NMDS and PERMANOVA, respectively.

## 3 RESULTS

### 3.1 Capture-based sampling (CBS) and literature survey

We performed a total of 16 surveys from March 2015 to November 2017, covering all seasons and months during the study period, with the exception of two winter months (January and February) and a summer month (July). In total, we recorded a total of 217 fish species distributed across 48 families and 128 genera. Results from CBS, supplemented by photographic observations of the highly mobile species, are summarized in Table S1.

A comprehensive literature survey across the atlas of Japanese fishes (Nakabo, 2013), which was supplemented by more recent information from the website (http://www.fish-isj.jp/info/list_additon.html), showed that 1,673 species belonging to 138 families and 586 genera are likely to occur in coastal waters of Okinawa Islands (Table S2).

### 3.2 eDNA metabarcoding (EDM)

The MiSeq paired-end sequencing (2 × 150 bp) of the 11 libraries, together with an additional 29 libraries (total = 40), yielded a total of 4,452,873 reads, with an average of 96.2% base calls, with Phred quality scores of ≥30.0 (Q30; error rate = 0.1% or base call accuracy = 99.9%). This run was highly successful considering the manufacture’s guidelines (Illumina Publication no. 770-2011-001 as of 27 May 2014) are >80% bases ≥Q30 at 2 × 150 bp.

Of the 4,452,873 reads, a total of 1,266,338 reads were assigned to the 11 libraries, and the numbers of raw reads for each library were relatively uniform, ranging from 89,750 to 141,322 with an average of 113,138 reads (Table 2). After merging two overlapping paired-end fastq files (1,244,520 reads [98.3%]), the primer-trimmed sequences were subjected to quality filtering to remove low-quality reads (1,234,627 reads [97.5%]). The remaining reads were dereplicated for subsequent analysis, and single-to tripletons were removed from the unique sequences. Then, reads were denoised to remove putatively erroneous and chimeric sequences, and the remaining 1,021,542 reads (80.7% of the raw reads) were subjected to taxon assignments after rarefaction to the minimum number of reads (i.e., 72,302). Of these, 999,296 reads (97.8% of the denoised reads) were putatively considered as fish sequences, and BLAST searches indicated that non-fish sequences (22,246 reads [2.2%]) mostly consisted of mammals (i.e., cows, pigs, and humans) and a few unknown sequences. The four negative controls (i.e., FB, EB, 1B, and 2B) were subjected to the same analysis pipeline and yielded only 99 denoised reads in total (Table 1; only 0.008% of the total raw reads), which were not taken into consideration in the subsequent analyses as their subtraction from the corresponding species did not affect the presence/absence data matrix.

**TABLE 2.** Summary of read numbers after data preprocessing (the second to fifth columns) and after taxon assignment (the last two columns) for the 11 libraries and four blanks. The numbers in parentheses in the third to fifth columns are percentages of the raw read numbers, while those in the last two columns are percentages of denoised read numbers. FB, EB, 1B, and 2B indicate blank samples from filtration, DNA extraction, 1st PCR, and 2nd PCR, respectively.

| Library | Raw read | Merged | Q filtered | Denoised | Fish | Non-fish |
| --- | --- | --- | --- | --- | --- | --- |
| Station 1 | 103,063 | 101,850 (98.8) | 101,197 (98.2) | 85,576 (83.0) | 85,445 (99.8) | 131 (0.2) |
| Station 2 | 137,257 | 135,779 (98.9) | 135,177 (98.5) | 115,590 (84.2) | 113,782 (98.4) | 1,808 (1.6) |
| Station 3 | 127,559 | 125,686 (98.5) | 124,698 (97.8) | 104,908 (82.2) | 100,889 (96.2) | 4,019 (3.8) |
| Station 4 | 139,776 | 136,938 (98.0) | 135,364 (96.8) | 113,505 (81.2) | 112,381 (99.0) | 1,124 (1.0) |
| Station 5 | 111,563 | 108,575 (97.3) | 107,531 (96.4) | 84,153 (75.4) | 80,180 (95.3) | 3,973 (4.7) |
| Station 6 | 104,622 | 102,356 (97.8) | 101,617 (97.1) | 84,919 (81.2) | 83,308 (98.1) | 1,611 (1.9) |
| Station 7 | 91,112 | 89,750 (98.5) | 89,041 (97.7) | 72,641 (79.7) | 70,512 (97.1) | 2,129 (2.9) |
| Station 8 | 93,339 | 90,974 (97.5) | 89,960 (96.4) | 72,302 (77.5) | 69,408 (96.0) | 2,894 (4.0) |
| Station 9 | 99,870 | 97,704 (97.8) | 96,638 (96.8) | 77,007 (77.1) | 75,495 (98.0) | 1,512 (2.0) |
| Station 10 | 142,972 | 141,322 (98.8) | 140,658 (98.4) | 117,373 (82.1) | 115,372 (98.3) | 2,001 (1.7) |
| Station 11 | 115,205 | 113,586 (98.6) | 112,746 (97.9) | 93,568 (81.2) | 92,524 (98.9) | 1,044 (1.1) |
| Total | 1,266,338 | 1,244,520 (98.3) | 1,234,627 (97.5) | 1,021,542 (80.7) | 999,296 (97.8) | 22,246 (2.2) |
| FB | 304 | 289 (95.1) | 285 (93.8) | 285 (93.8) | 53 (100) | 0 (0.0) |
| EB | 279 | 272 (97.5) | 271 (97.1) | 271 (97.1) | 46 (100) | 0 (0.0) |
| 1B | 47 | 36 (76.6) | 32 (68.1) | 32 (68.1) | 0 (100) | 0 (0.0) |
| 2B | 0 | 0 | 0 | 0 | 0 | 0 |
| Total | 630 | 597 (94.8) | 588 (93.3) | 588 (93.3) | 99 (16.8) | 0 (0.0) |

After the automatic taxon assignments, we excluded deep-sea and pure freshwater fishes from the list as our major purpose was the detection of tropical and subtropical coastal fishes inhabiting the target coral reef lagoon, and not exogenous eDNA from adjacent ecosystems. After excluding such putatively exogenous eDNA, we visually inspected family level NJ trees and revised the species names in the list. The final list included a total of 886 detections, which were assigned to 291 species distributed across 49 families and 142 genera (Table S3). The number of species detected at each station ranged from 58 to 92, with a mean of 80.1.

Frequently detected species (≥ 6 detections, i.e., species detected by eDNA from more than half of the 11 stations) occupied nearly half of the total detections (432 [48.8%]), consisting of 54 species (18.3%) distributed across 18 families (35.2%) and 36 genera (25.7%). These frequently detected species were listed in Table 2 with frequency of occurrences, read abundances, and presence or absence of specimens from CBS.

### 3.3 Species richness

In total, 410 fish species were detected by considering both methods, i.e., 217 and 291 species by CBS and EDM, respectively, (Figure 2), and 98 species were detected by both (23.9% of the total detections). All species were distributed across 61 families and 201 genera, and both methods detected common genera and families at higher proportions than species-level detections (Figure 2; 69 genera and 35 families at 32.9.4% and 56.5%, respectively,). Of the 119 species solely detected by CBS, 96 species (81%) were found in the reference database.

**FIGURE 2.**
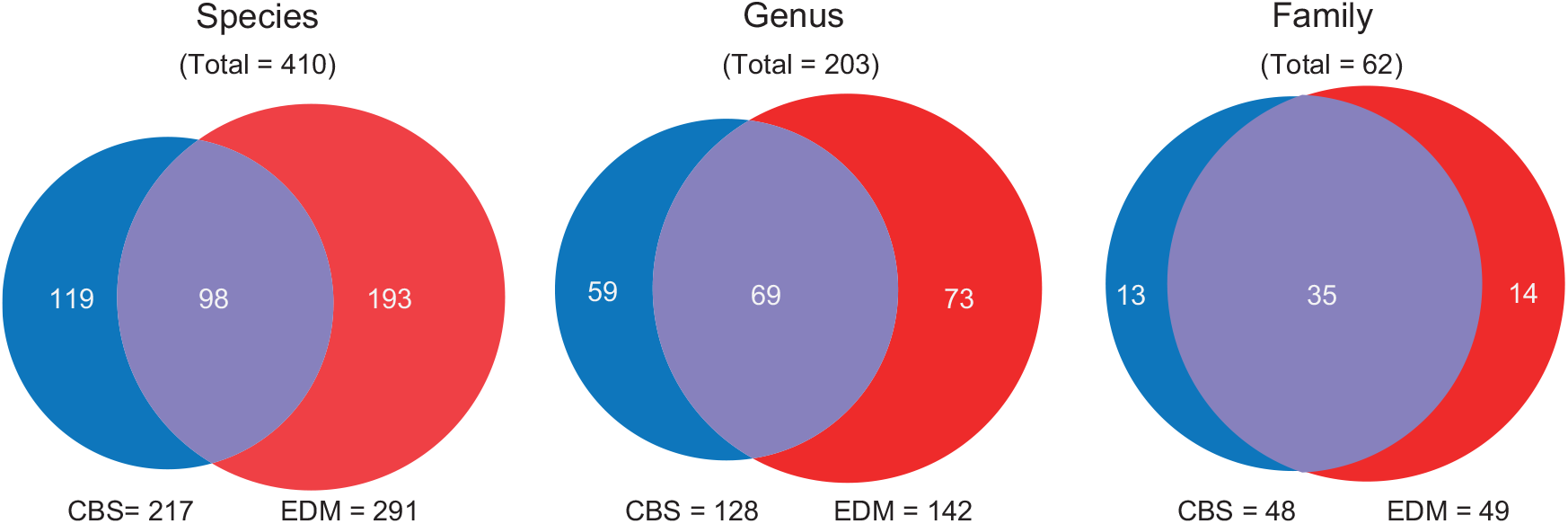
Venn diagrams of the number of detected fish species, genus, and family in capture-based sampling (CBS) and eDNA metabarcoding (EDM)

On the basis of the EDM data, *S*_max_ using the Chao II bias-corrected estimator and the species accumulation curve were calculated (Figure 3). EDM detected 291 species, with a calculated *S*_max_ of 407 (95% CI: 392-421) and 411 species by the Chao II estimator and species accumulation curve, respectively. Species accumulation curves suggested that at the 36 sampling sites of eDNA, *S*_max_ would reach that of Chao II as well as the total observed species by both methods (Figure 3). We found that species richness estimated from our data, and from the literature survey plus an additional two check lists (Senou, Kodato, Nomura, & Yunokawa, 2006; Senou, Kobayashi, & Kobayashi, 2007) consistently exhibit higher species richness than data from the visual transect censuses for the respective geographic scales (Figure 4).

**FIGURE 3.**
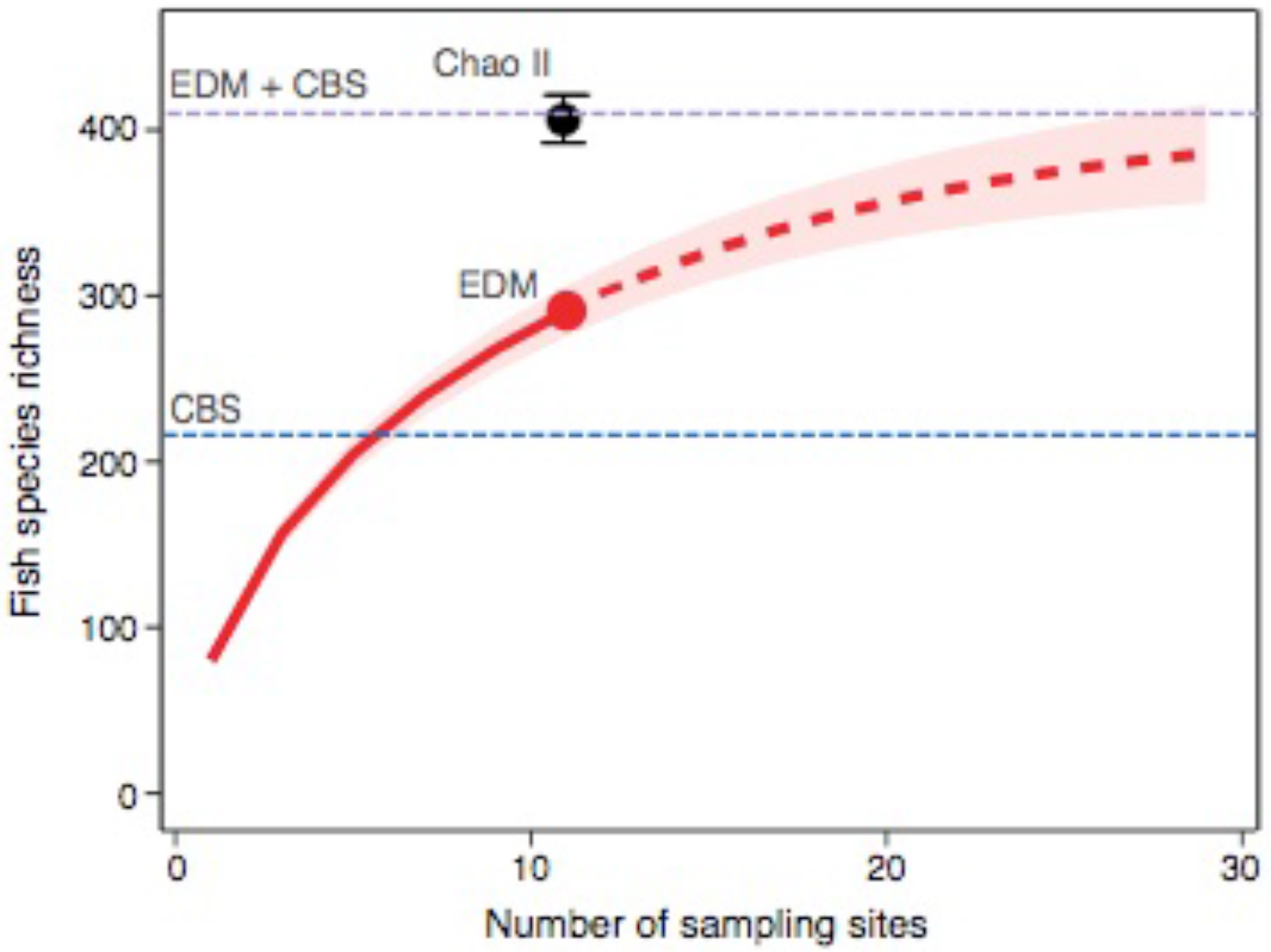
Species accumulation curves with 95% CI of the fish community as detected from eDNA metabarcoding (EDM) from the 11 sites (dotted line indicates extrapolated plots). The blue and purple dotted lines indicate the observed species number by capture-based sampling (CBS) and the combined species number from CBS and EDM, respectively. The black dot indicates the Chao II estimator (mean ± 95% CI)

**FIGURE 4.**
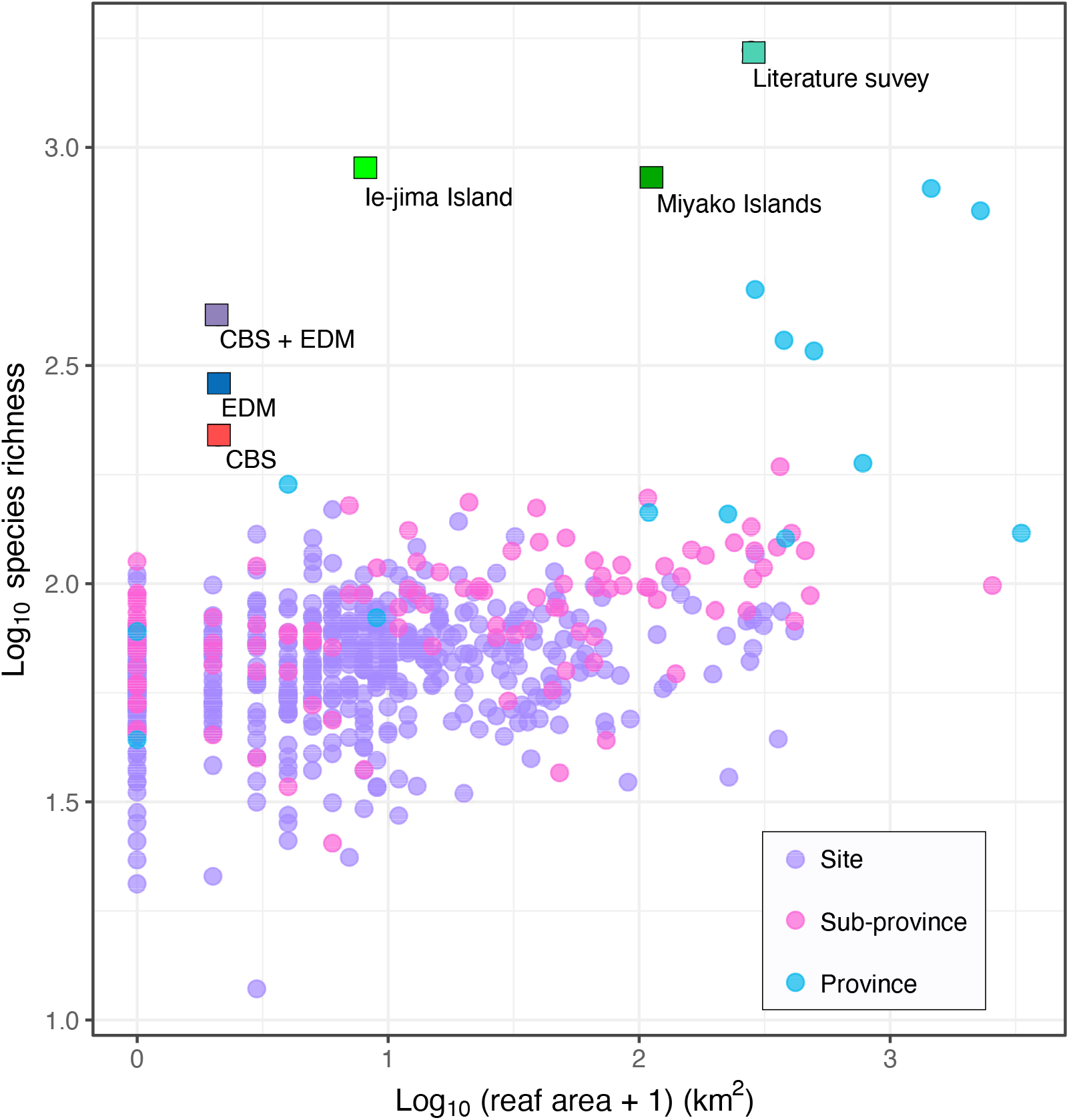
Relationships between the reef area and fish species richness taken from visual transect censuses from tropical reefs around the globe (Barneche et al. 2018, circle symbols). The data from 13,050 transects were assembled and transformed into species richness for 485 sites in 109 sub-provinces spread across 14 provinces by Barneche et al. (2018). Our data from capture-based sampling (CBS = 217 spp.), eDNA metabarcoding (EDM = 291 spp.), CBS + EDM (410 spp.) in the lagoon (1.1 km^2^), data resulting from the literature survey (1678 spp.; summed lagoon area = 278 km^2^), checklists of fishes from Ie-jima Island (Senou, Kobayashi, & Kobayashi, 2007) and from Miyako Island (Senou, Kodato, Nomura, & Yunokawa, 2006) are plotted on the figure as square symbols. Reef areas of the latter two islands were calculated using a function in Google Map (https://www.google.co.jp/maps)

### 3.4 Fish community structure

NMDS clearly showed dissimilarity in fish community composition between the four sites along the offshore reef edge (Sts. 5-8) and the seven on shore-side seagrass beds sites (Sts. 1-4 and 9-11) (Figure 5, NMDS stress = 0.139). PERMANOVA detected significant differences in community composition between the offshore and shore-side sites (*F* = 1.54, *P* = 0.012, *N* = 11).

**FIGURE 5.**
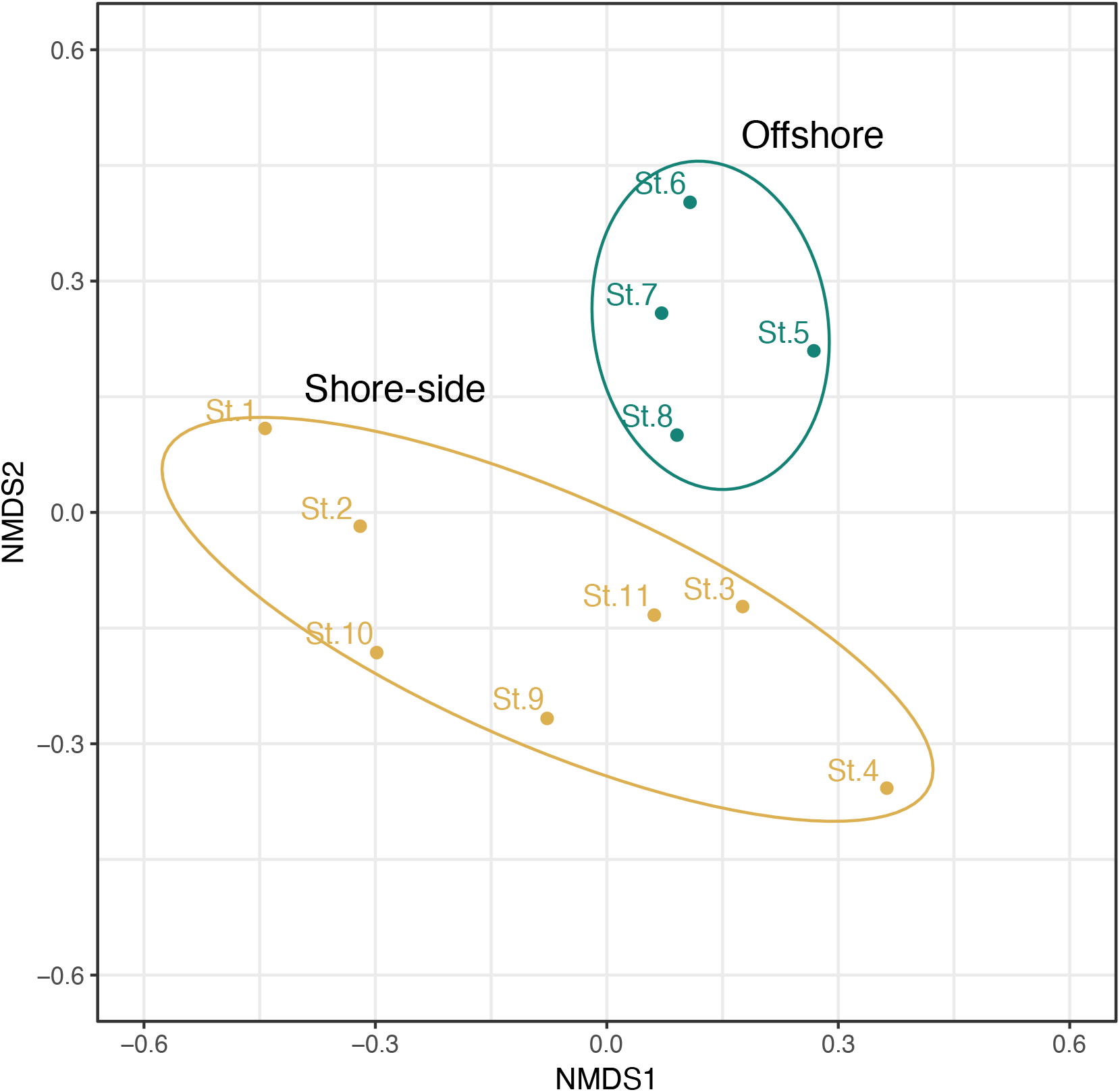
Non-metric multi-dimensional scaling (NMDS) for fish communities detected at four sites along the offshore reef edge (Sts. 5-8) and seven sites on shore-side seagrass beds (Sts. 1-4 and 9-11). NMDS stress was 0.139

### 3.5 Comparison of ecological and biological traits

We compared the fish ecological/biological traits between frequently detected species (i.e., with more than six detections; Table 3) along the offshore reef-edge sites (Sts. 5-8) and on the shore-side seagrass-bed sites (Sts. 1-4 and 9-11) (Figure 6). GLM found a significant difference in the maximum depth range to deep (m) between the offshore reef edge and the shore-side seagrass beds (Figure 6E; Table 6).

**TABLE 3.**
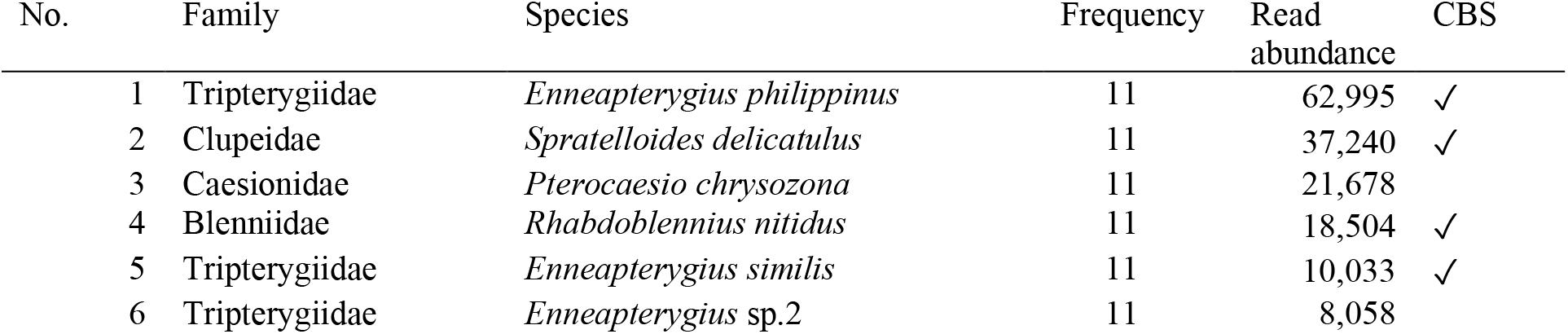
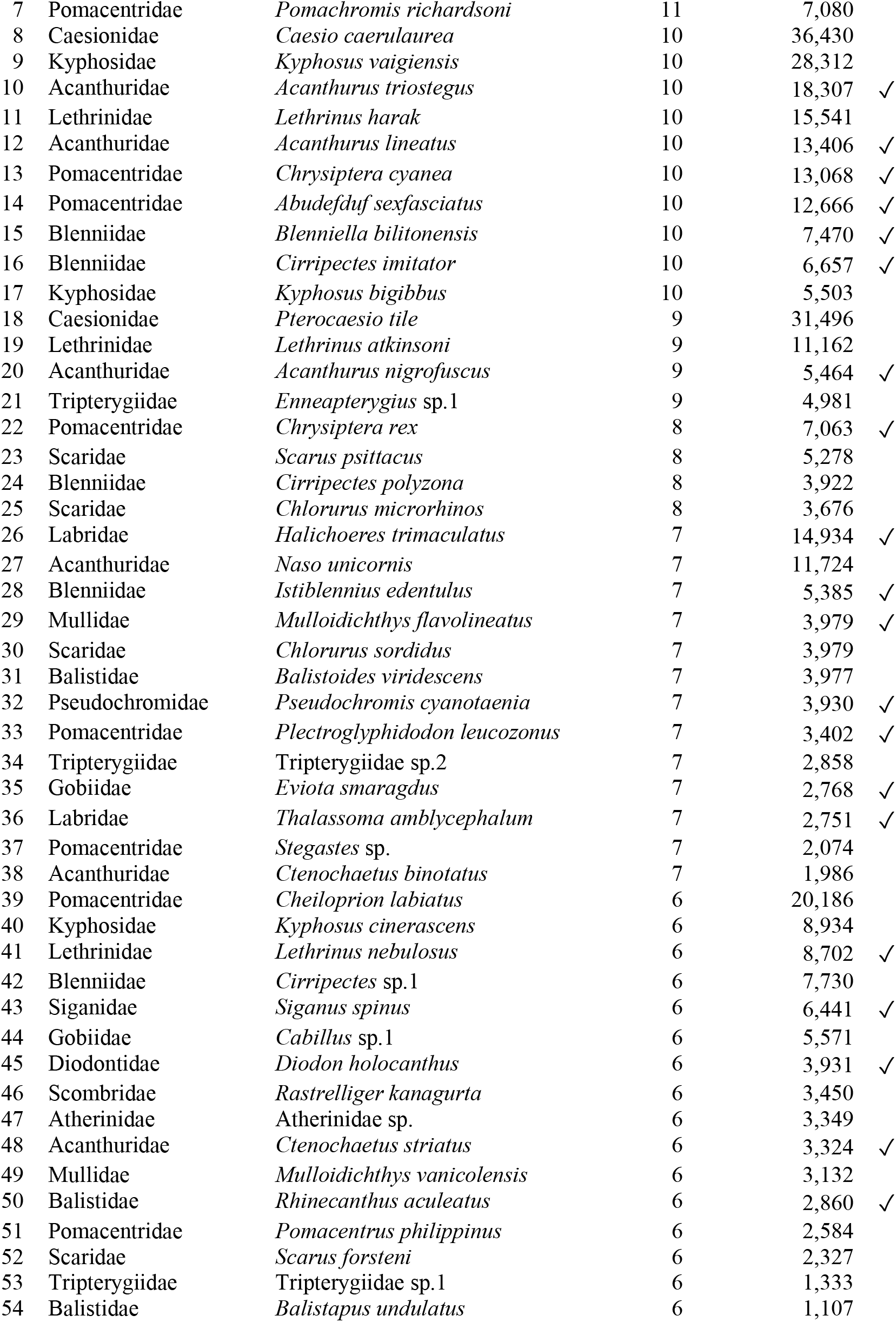
List of fish species with six or more eDNA detections from the 11 stations in a coral reef lagoon off Bise, Okinawa, southern Japan. Fish species were arranged in descending order of detection frequencies followed by read abundances. Species recorded in capture-based sampling (CBS) were checked in the rightmost column

**FIGURE 6.**
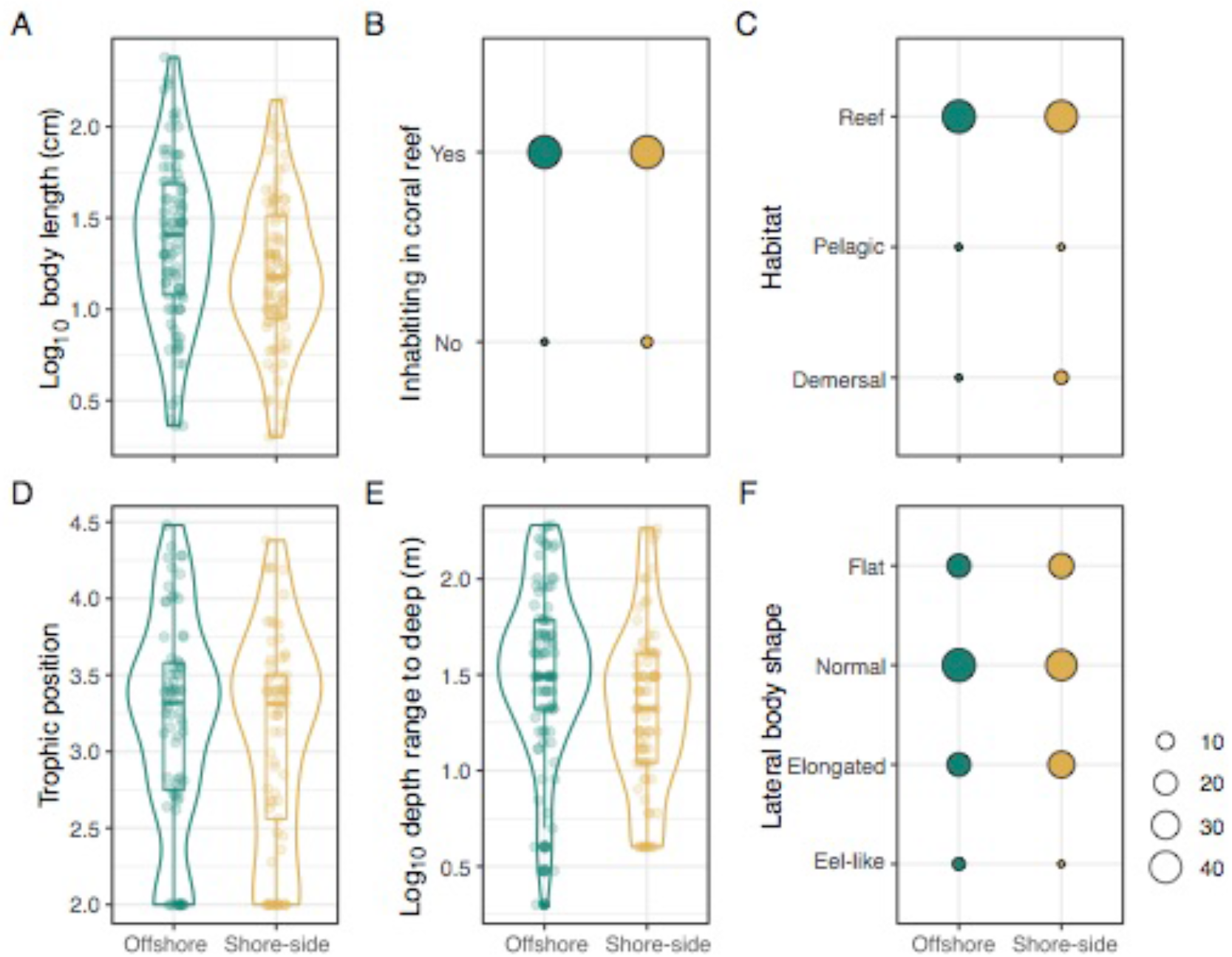
Comparisons of the six biological and ecological fish traits between the frequently detected species along the offshore reef edge (Sts. 5-8) and on the shore-side seagrass beds (Sts. 1-4 and 9-11) evaluated by indicator taxa analysis (Tables 5 and S5). The traits include A) body length (cm), B) habitat preference in the coral reefs, C) habitat types, D) trophic positions, E) depth range to the deep, and F) lateral body shape types. Boxes and bars in the box plots indicate median ± inter-quartiles and ± 1.5 × inter-quartiles. The smooth peripheral lines indicate the distributions of the data using a violin plot. The points represent each data value. In the balloon plots, balloon sizes are proportional to the number of species in each category, as indicated in the legend.

## 4 DISCUSSION

### 4.1 Species richness

Estimating the species richness in a coral reef lagoon by traditional capture-based sampling (CBS) and/or visual observation presents a number of difficulties. Common resident species in the lagoon may be easier to directly capture or visually observe than migrant, nocturnal, or relatively rare species. In addition, corals create complex and microscale habitats, with many undescribed small and elusive species (Reimer et al., 2019). Seagrass beds also offer hiding places for several fishes and physically prevent their visual observation and CBS. Furthermore, several coral reef fishes have fossorial life styles and some even demonstrate mimicry (Nakabo, 2013), making them difficult to capture and visually observe. In addition to these biological and environmental factors, significantly high species diversity precludes accurate estimation of species richness. Sanciangco, Carpenter, Etnoyer, & Moretzsohn (2013) recently demonstrated that more than 2,000 fish species are likely to occur in the “Coral Triangle,” a diversity hotspot in the Indo-Pacific region, and its northern edge corresponds to the sampling sites of this study. Literature survey results (Table S2) were consistent with their estimation, listing 1,673 coastal fishes as likely to occur in the surrounding waters of the Okinawa Islands.

If all (or most) fish species release DNA into environmental water and their fragments can be co-amplified by the universal PCR primers, simultaneous and parallel detection of multiple species using eDNA metabarcoding (EDM) could avoid many of the difficulties associated with fish biodiversity monitoring by traditional methods (Thomsen et al., 2012). In fact, while only 217 fish species were confirmed in the 16 capture-based surveys over a two-year period (2015-2017), 291 species were detected by the subsequent EDM from 11 sites in just one hour per day of the survey in May 2019. Combining these two different approaches to biodiversity monitoring results in 410 detected fish species (Figure 2) distributed across 119 families and 193 genera, nearly doubling those confirmed by CBS. Of these 410 species, only 96 (24% of the total) were commonly identified by both methods (Figure 2), indicating that CBS failed to collect many of the species (195 spp.) detected by EDM. However, EDM did not detect 119 species confirmed by CBS (Figure 2). Nevertheless, given that the latter was based on a total of 16 surveys over two years covering nine months (Table S1), this gap may be filled by additional eDNA surveys in the future. Actually, extrapolation of the species accumulation curve (Figure 3, Table 4) suggested that an additional three or more surveys for the 11 sites (36 samples) would detect 90% of the 410 species, which would be worth corroborating in future studies.

**TABLE 4.**
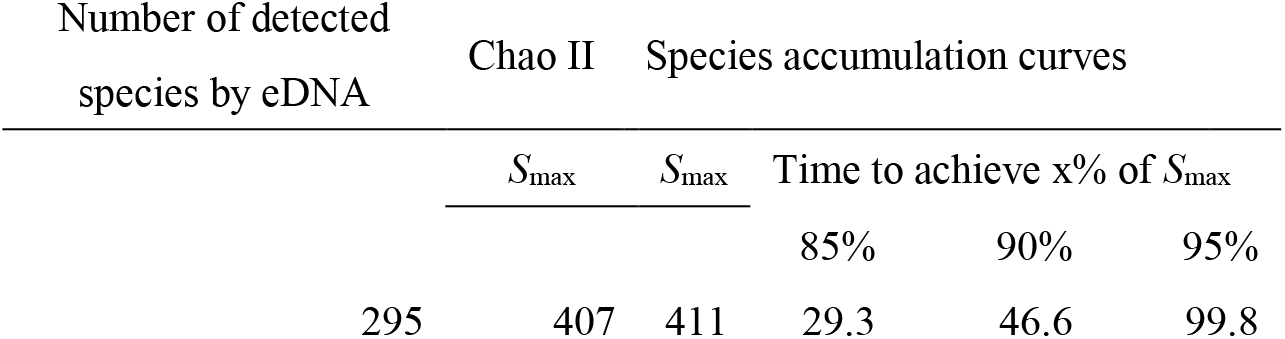
Summary of species richness estimation from expected total number (*S*_max_) using the Chao II bias-corrected estimator (Chao, 2005) and species accumulation curve model based on eDNA metabarcoding data

Remarkably, the 410 fish species identified by the two different methods (CBS and EDM) were close to the maximum numbers estimated by the Chao II estimator and species accumulation curve (407 and 411 species, respectively). Both estimations were based on EDM (291 species detected from 11 sites), suggesting that EDM is useful to estimate species richness, even in an ecosystem with remarkable species diversity, such as the coral reef lagoon. However, as stated above, our literature survey indicated that 1,673 coastal species are likely to occur in the surrounding waters of Okinawa Islands where the lagoon is located. This appears to be far from the 410 species identified by the two methods in the lagoon. However, it may be a reasonable figure considering the total lagoon area on the main island of Okinawa (278 km^2^; Fujiwara, 1994) and that of the lagoon in Bise (1.1 km^2^). In fact, the logarithms of the area and the number of species exhibit a clear positive proportional relationship for a wide variety of organisms and ecosystems at various geographic scales (Lomolino, 2000).

Although we have no comparable estimations of species richness at different geographic scales on Okinawa Island, visual transect censuses of reef-fish species have been conducted on a global scale (Barneche et al., 2018), and it would be meaningful to compare the number of species observed through such censuses with the present estimations. Recently, Barneche et al. (2018) compiled a database encompassing 13,050 visual transect censuses of reef-fish species and investigated biotic and abiotic factors responsible for changes in species richness at three geographic scales in 485 sites of 109 sub-provinces spread across 14 biogeographic provinces. Based on analysis using hierarchical linear Bayesian models, Barneche et al. (2018) found that reef area is one of the major predictors of species richness and that increasing habitat area yields higher species richness across the three geographic scales with different corresponding parameters.

As noted by Barneche et al. (2018), species richness increased with reef area, but with different slopes at different geographic scales. However, that our six data are plotted at upper outlier positions from those taken by the visual observations is quite reasonable, as observers are likely to overlook elusive or rare species during visual transect censuses, and the list of species based on visual observations tends to mostly consist of common and abundant species, resulting in underestimation of the species richness (Edgar et al., 2004; Harvey et al., 2004). Interestingly, the total number of species observed in this study (410 spp.), which is in good agreement with two estimations of species richness based solely on EDM from the 11 stations (407 and 417 spp.; see Table 4), and that of the literature survey (1,673 spp.) were 4.14 and 3.54 times higher, respectively, than those taken from census data with the same areas (99 and 472 spp. from 1 km^2^ from the Mexican Caribbean and 289 km^2^ from the Central Pacific, respectively). Further empirical studies are needed to address the validity of species-richness estimation based on EDM and the species-area relationships at different geographic scales in coral reef lagoons.

### 4.2 Habitat segregation

In the coral reef lagoons of Okinawa Islands, coral assemblages dominated along the offshore reef edge, with a strong wave action at high tide, while seagrass beds dominated along shore-side areas, with a calm wave condition throughout the year. These two adjacent areas form a coral reef-seagrass gradient (Dorenbosch et al., 2005). In the case of the coral reef lagoon in Bise (Figure 1), Sts. 5-8 were located along the reef edge and there were patchy coral assemblages, mainly consisting of acroporid corals on the sandy bottom. Meanwhile, eelgrass beds spread across shore-side sites (Sts. 1-4 and 9-11), basically on sandy bottoms with little sediment and small coral assemblages scattered at some sites (Sts. 9-11). During seawater sampling on May 8, 2019, the reef edge was submerged underwater at high tide (ca. 180 cm above the lowest tide level; see Figure 1E), and open ocean swells crossed the reef edge and entered the lagoon, even making the small-boat navigation somewhat difficult around Sts. 5-8. Contrastingly, the swells subsided toward the coastline, and the sea was calm in the shore-side sites around Sts. 1-4 and 9-11.

These environmental gradients produce different habitats, resulting in different fish community compositions comprising a mixture of reef- and seagrass-associated species, nursery species, generalists, and rare species (Dorenbosch et al., 2005). In this study, however, we were unsure *a priori* about the detectability of eDNA for differentiating the two fish communities in different habitats within the small lagoon. In fact, the tidal difference was large (approximately 2 m), and offshore water flowed into the shore-side sites on the ebb tide toward a high tide time (08:43) during sampling (07:40-08:50). This apparent inflow from the reef edge should have created outflowing compensation currents, supposedly resulting in a complex mixture of eDNA from different habitats. Furthermore, the average distance from the offshore sites and the corresponding nearest shore-side sites was approximately 350 m, which is close to the marginal distance (300 m) that eDNA can reach in the coastal inner bay with low tidal differences (< 30 cm) and no strong currents across the bay (Murakami et al., 2019).

Nonetheless, NMDS clearly showed dissimilarity in the detected fish communities between the four sites along the offshore reef edge (Sts. 5-8) and seven sites on shore-side seagrass beds (Sts. 1-4 and 9-11) (Figure 5). In addition, PERMANOVA detected statistically significant differences in community compositions between the offshore and shore-side sites. We also compared the six biological and ecological traits between the frequently detected species (≥ 6 detections) along the offshore reef edge (Sts. 5-8) and on the shore-side seagrass beds (Sts. 1-4 and 9-11) (Figure 6). The maximum depth range to deep (m) was the only trait that exhibited a statistically significant difference (Figure 6E) and would be a critical trait for distributing in the offshore reef edge, while no such differences were found for the other traits, including body length and types, trophic position, and habitat preferences (Figure 6). Andradi-Brown et al. (2018) also found a significant depth gradient in reef-fish community changes with the traditional SCUBA survey.

To see what species actually contribute to such distinct community differences, we divided the list of species from EDM (Table S3) into the following three categories by detection frequencies (Table S4), i.e., those occurring 1) uniquely to the offshore reef-edge sites (Sts. 5-8); 2) uniquely to the shore-side seagrass-bed sites (Sts. 1-4 and 9-11); and 3) across both regions (67, 110, and 114 spp. respectively). Of the four sites along the offshore reef edge, *Parupeneus multifasciatus* Quoy Gaimard (Manybar goatfish) were found at all four sites, followed by *Centropyge ferrugata* Randall & Burgess (Rusty angelfish), *Eviota guttata* Lachner & Karnella (Spotted dwarf goby), *Platybelone argalus platyura* Bennett (Keeled needlefish), *Thalassoma lutescens* Lay & Bennett (Yellow-brown wrasse), and *Scarus schlegeli* Bleeker (Yellowband parrotfish), which were found at the three sites. Of the seven sites on the shore-side seagrass beds, Atherinidae sp. (a species of silversides), *Pomacentrus philippinus* Evermann & Seale (Philippine damsel), and *Rastrelliger kanagurta* Cuvier (Indian mackerel) were uniquely found in the six sites. Results from the indicator taxa analysis are congruent with the above nine unique and frequently detected species in the two respective regions, with an additional *Mellichthys* sp.1 (a species of pinktailed triggerfish) as a representative of the offshore sites (Table 5). The latter species was detected from all four offshore reef-edge sites, but it was also detected from a single site (St. 10) on the shore-side seagrass-bed sites. With the exception of *R. kanagurta*, highly significant detections of the remaining nine species in either region are congruent with our empirical observations in the lagoon. However, *R. kanagurta* is highly pelagic, often found outside the lagoon. As the eDNA survey was conducted in May, during its spawning season (April to August; Uehara, Motonaga, Tachihara, Ohta, & Ebisawa, 2015), we may have incidentally detected a patchy stream of its gametes in the shore-side sites on seagrass beds.

**TABLE 5.** Significantly highly-frequently detected fish taxa either in the offshore reef-edge sites (Sts. 5-8) or in the shore-side seagrass-bed sites (Sts. 1-4 and 9-11). Best represents the most frequently detected sites. *P* value was calculated with 999 permutations after Sidak’s correction for multiple testing. Statistical data for those species other than listed below are shown in Table S5

| Species | Offshore | Shore-side | Best | $P$ value |
| --- | --- | --- | --- | --- |
| *Parupeneus multifasciatus* | 0.003 | 1 | Offshore | 0.005991 |
| *Atherinidae* sp. 1 | 1 | 0.011 | Shore side | 0.021879 |
| *Pomacentrus philippinus* | 1 | 0.014 | Shore-side | 0.027804 |
| *Rastrelliger kanagurta* | 1 | 0.014 | Shore-side | 0.027804 |
| *Centropyge ferrugata* | 0.016 | 1 | Offshore | 0.031744 |
| *Mellichthys* sp. 1 | 0.018 | 1 | Offshore | 0.035676 |
| *Eviota guttata* | 0.021 | 1 | Offshore | 0.041559 |
| *Platybelone argalus platyura* | 0.023 | 1 | Offshore | 0.045471 |
| *Thalassoma lutescens* | 0.023 | 1 | Offshore | 0.045471 |
| *Scarus schlegeli* | 0.023 | 1 | Offshore | 0.045471 |

**TABLE 6.** Results of generalized linear model for Figure 6. Bold font indicates a significant coefficient

| Items | Coefficient | Std. error | z-value | p-value |
| --- | --- | --- | --- | --- |
| (Intercept) | 1.9448 | 2.3505 | 0.827 | 0.408 |
| $\log_{10}$ body length | -0.7016 | 0.6422 | -1.092 | 0.2746 |
| Inhabiting in coral reef | -0.5248 | 0.6962 | -0.754 | 0.451 |
| Habitat | -0.8685 | 2.0647 | -0.421 | 0.674 |
| Trophic position | 0.2934 | 0.3133 | 0.937 | 0.3489 |
| **$\log_{10}$ depth range to deep** | **-1.2308** | **0.4931** | **-2.496** | **0.0126** |
| Lateral body shape | 0.7297 | 1.0783 | 0.677 | 0.4986 |

Seagrass beds are known to harbor a number of unique resident species compared with adjacent coral reefs (Nakamura & Tsuchiya, 2008), as coral reef resident species occasionally use seagrass beds as nursery or foraging habitats, while seagrass-bed resident species generally do not use coral reefs (Horinouchi, Nakamura, Sano, & Shibuno, 2005; Nagelkerken et al., 2000). Accordingly, a list of seagrass-bed resident species would be useful to characterize the fish communities in the lagoon. For conservation of resident fishes in seagrass beds in Sekisei Lagoon, approximately 400 km southwest of Okinawa Island, Horinouchi et al. (2005) conducted visual transect censuses to reveal fish community structures in six seagrass beds and adjacent coral reefs in the lagoon. They observed 46 fish species across the six seagrass beds, with 30 species being restricted to seagrass beds and not found in the adjacent coral reefs. Among these 30 species, nine species were also found in our list (Table S4), and out of these, four species were restricted to the shore-side seagrass-bed sites in this study, i.e., *Parupeneus indicus* Shaw (Indian goatfish), *Novaculoides macrolepidotus* (Seagrass wrasse), *Siganus argenteus* Quoy & Gaimard (Streamlined spinefoot), and *Siganus fuscescens* Houttuyn (Dusky spinefoot). Meanwhile, *Parupeneus barberinoides* Bleeker (Bicolor goatfish) and *Parupeneus ciliatus* Lacepède (Whitesaddle goatfish) were found only along the offshore reef-edge sites, while *Lethrinus harak* Forsskål (Thumbprint emperor), *Lethrinus atkinsoni* Seale (Atkinson’s emperor), and *Lethrinus nebulosus* Forsskal (Spangled emperor) were found in both sites. Therefore, the fish community data from the seagrass beds in Sekisei Lagoon (Horinouchi et al., 2005), did not fully characterize the seagrass bed community in this study.

## 5 CONCLUSIONS

This study demonstrates that EDM is a rapid, cost-effective, and useful approach for estimating species richness in tropical and subtropical fish communities with remarkable species diversity. Additionally, it also showed that it was not possible to comprehensively cover those fish species with a limited number of samples (11 in this study) even in a small lagoon (1.1 km^2^), and future repetitive sampling would be required to verify the predicted species richness (around 410 spp.) in the lagoon. In addition to the species richness estimation, data from EDM could detect a known habitat segregation of fish communities between the offshore reef edge and shore-side seagrass beds. However, there is ambiguity as to whether eDNA data reflected true fish communities in the lagoon. Thus, concurrent implementation of CBS/visual observations and eDNA surveys would help to resolve such ambiguity concerning the detectability of EDM.

## ACKNOWLEDGEMENTS

We thank K. Ueda and K. Sato for supporting this study. R. Nozu helped us perform the statistical analysis. We also thank D. Barneche for kindly providing his compiled data with us for Figure 4. This work was supported by Japan Science and Technology Agency CREST Grant Number JPMJCR13A2, Japan Society for the Promotion of Science KAKENHI Grant Number JP19H03291, and Ministry of Education, Culture, Sports, Science and Technology, the Ocean Resource Use Promotion Technology Development Program Grant Number JPMXD0618068274.

## AUTHOR CONTRIBUTION

SO and MM designed the study and performed eDNA sampling. SO., KM, and NH conducted the traditional capture-based sampling and literature survey. TS and MM conducted the laboratory work and bioinformatic analysis of the NGS data. HD and SO conducted the statistical analyses. MM, SO, and HD wrote the manuscript with all authors giving final approval for publication.

## DATA ARCHIVING STATEMENT

The minimal raw dataset is uploaded to the DDBJ Sequence Read Archive (https://www.ddbj.nig.ac.jp/dra/index-e.html; Accession number: DRA009512).

